# Action of cofilin on ADP-Pi-containing actin filaments

**DOI:** 10.64898/2026.09.11.751010

**Authors:** Inaara Kassamaly, Natalia Nojszewska, Thomas Dos Santos, Hugo Wioland, Antoine Jégou, Guillaume Romet-Lemonne

**Affiliations:** Université Paris Cité, CNRS, Institut Jacques Monod, 75013 Paris, France

## Abstract

In cells, the disassembly of actin filaments is tightly controlled by molecular mechanisms where cofilin plays a central role. Cofilin binds to actin filaments cooperatively, forming clusters that lead to the severing of the filaments. Since cofilin binds preferentially filaments that have hydrolysed ATP and released inorganic phosphate (P_i_), most studies have focused on the action of cofilin on ADP-actin filaments. Here, we ask how cofilin binds and severs filaments that still contain a large fraction of ADPꞏP_i_-actin. To answer this, we use microfluidics and fluorescence microscopy to monitor single filaments exposed to cofilin in controlled conditions, allowing us to disentangle and to quantify the different reactions. We find that the step limiting cluster nucleation is the binding of 2 cofilins on a minimal group of 3-4 contiguous ADP-actin subunits. Cofilin clusters then expand mostly in the direction of the pointed end of the filament, by easily binding to the ADP-actin neighbor of an ADPꞏP_i_- actin subunit, whose P_i_ release it accelerates. Severing at the ADPꞏP_i_-actin/cofilin cluster boundary is 30-fold slower than for an ADP-actin boundary. Globally, it seems that cofilin senses and modifies the nucleotide state of the actin subunits that are in its direct vicinity only. Our results provide a more comprehensive understanding of the action of cofilin, taking into account the nucleotide-state complexity of actin filaments.

## Introduction

The actin cytoskeleton is involved in a number of essential cellular processes, including cell motility, cell division and endocytosis (*1*). To perform these tasks, actin monomers assemble into filaments which are polar, with a barbed end (BE) and a pointed end (PE). Actin filaments are organized in higher order networks which can generate mechanical forces, and eventually disassemble back to the monomeric state.

This turnover of actin, from monomeric (G-actin) to filamentous (F-actin) form and back, is an essential feature. It is linked to the nucleotide state of actin. G-actin in the cytoplasm is predominantly loaded with ATP, which is hydrolyzed within seconds into ADP + P_i_ (inorganic phosphate) after incorporation at the barbed end of the filament (*2*, *3*). ADP stays locked inside F-actin as the P_i_ is released, with a half-life of approximately 100 seconds (*4–7*). It is only when actin has depolymerized back to G-actin that its ADP can be exchanged for ATP from the cytoplasm.

In cells, the different steps of actin turnover need to be precisely controlled. The disassembly of actin filaments, in particular, can take place at vastly different rates depending on the networks and their cellular context (*8*). Several proteins can take part in regulating the disassembly of actin filaments, and proteins of the ADF/cofilin family play a central role in accelerating disassembly (*9*).

Thanks to decades of research from several fields (*10–13*), the action of cofilin on actin filaments can be described as follows. Cofilin binds to the sides of ADP-actin filaments, where one cofilin bridges two longitudinal actin neighbors. This binding is cooperative, favoring the addition of cofilin near sites that are already occupied by cofilin (*14*). This results in the formation of cofilin clusters, which locally alter the conformation of the actin filament. As a consequence, thermal fluctuations cause filaments to sever at the boundaries of cofilin clusters. This can be understood as resulting from the local weakening of actin-actin bonds due to their cofilin-induced conformational change (*12*, *15*, *16*). Alternatively, severing has been proposed to result from the mechanical mismatch between the stiffer, bare F-actin region and the softer cofilin-decorated region (*17*, *18*). Cofilin-saturated filaments do not sever, but depolymerize from both ends, barbed end included, even in the presence of G-actin and capping protein (*19*, *20*). We now have a good mechanistic understanding of these different events. Individual reaction rates have been measured, and high-resolution cryo-EM data have revealed the conformational changes underpinning these mechanisms (*21*).

There are still, however, some open questions. How cofilin clusters nucleate, and in particular what would be the minimal number of cofilins required for cooperative binding to be effective, is unclear. Structural data indicate that this minimal number could be as low as two (*22*), but kinetic data and mechanistic insights are lacking. The distance over which cooperativity is effective (range) is also an unsettled question. CryoEM data indicate that the conformational changes on actin that result from the binding of cofilin do not expand beyond the nearest neighbor (*16*, *22*), but measurements on dynamic filaments using high-speed AFM (*23*) or single molecule fluorescence microscopy (*24*) detect structural modifications and enhanced cofilin recruitment several subunits away from the nearest cofilin. Finally, while there seems to be a consensus on the asymmetry of severing, found by several groups to be more likely to occur at the PE-side boundary of cofilin clusters (*19*, *25*, *26*), contradicting results have been published on cluster growth, with some studies reporting a faster expansion towards the PE (*23*) and others reporting equal velocities in both directions (*19*, *20*).

The largest gap in our understanding concerns the action of cofilin on freshly assembled, ADPꞏPi-rich actin filament regions. This question bears importance in the cell context, where actin filaments are exposed to cofilin as they grow, and are thus made of freshly assembled ADPꞏP_i_-actin, and where cytosolic P_i_ concentrations are likely to maintain a fraction of F-actin in the ADPꞏP_i_ state (*27*, *28*). Yet, nearly all the available mechanistic results, whether kinetic rates or structural details, are for cofilin binding to ADP-actin filaments.

Early studies in bulk solution have established that cofilin binds preferentially to ADP-actin (*29*, *30*), and recent cryoEM data provide structural explanations for this preference (*31*). It has also been established early on that cofilin accelerates the release of Pi from ADPꞏP_i_-actin filaments (*30*). The generally accepted mechanism is that cofilin bound to ADP-actin favors the release of Pi from nearby ADPꞏP_i_-actin subunits. Based on this model, Suarez et al. have analyzed images of single filaments decorated by fluorescently labeled cofilin and concluded that Pi release is accelerated by one order of magnitude, up to tens of actin subunits away from a cofilin cluster boundary (*32*). This distance seems at odds with the cryo-EM data that indicate that the conformational changes of actin induced by the binding of cofilin are very local (*16*, *22*). To our knowledge, no other kinetic measurements have addressed this question since, and how cofilin clusters expand in a landscape of mixed nucleotides remains unclear. Most other aspects of the interaction of cofilin and ADPꞏP_i_-actin subunits are unexplored. There is no model for how clusters are nucleated on filaments that contain a mix of ADP- and ADPꞏP_i_-actin subunits. No structural data are available for ADPꞏP_i_-actin/cofilin cluster boundaries, and we do not know if they are less stable than ADP-actin/cofilin cluster boundaries.

To answer these questions, we monitored single actin filaments exposed to cofilin, in different situations. This allowed us to disentangle the multiple reactions that can take place, and to shed light on the nucleation of cofilin clusters, their growth, and their ability to sever actin filaments in an ADPꞏP_i_- rich context. Doing so has also provided new insights into the action of cofilin on ADP-actin filaments.

## Results

### Microfluidics allows to monitor individual reactions on actin filaments

To disentangle the many reactions that take place when cofilin decorates and severs actin filaments (Fig. 1a) we have taken advantage of microfluidics for the study of single actin filaments using wide field and TIRF microscopy (*4*, *33*). This technique has already proven itself as a valuable tool to monitor individual reactions on ADP-actin and quantify their rates (*19*). In particular, it allows to rapidly change the solution to which filaments are exposed, and thus allows to create conditions where specific reactions can be quantified.

**FIGURE 1.**
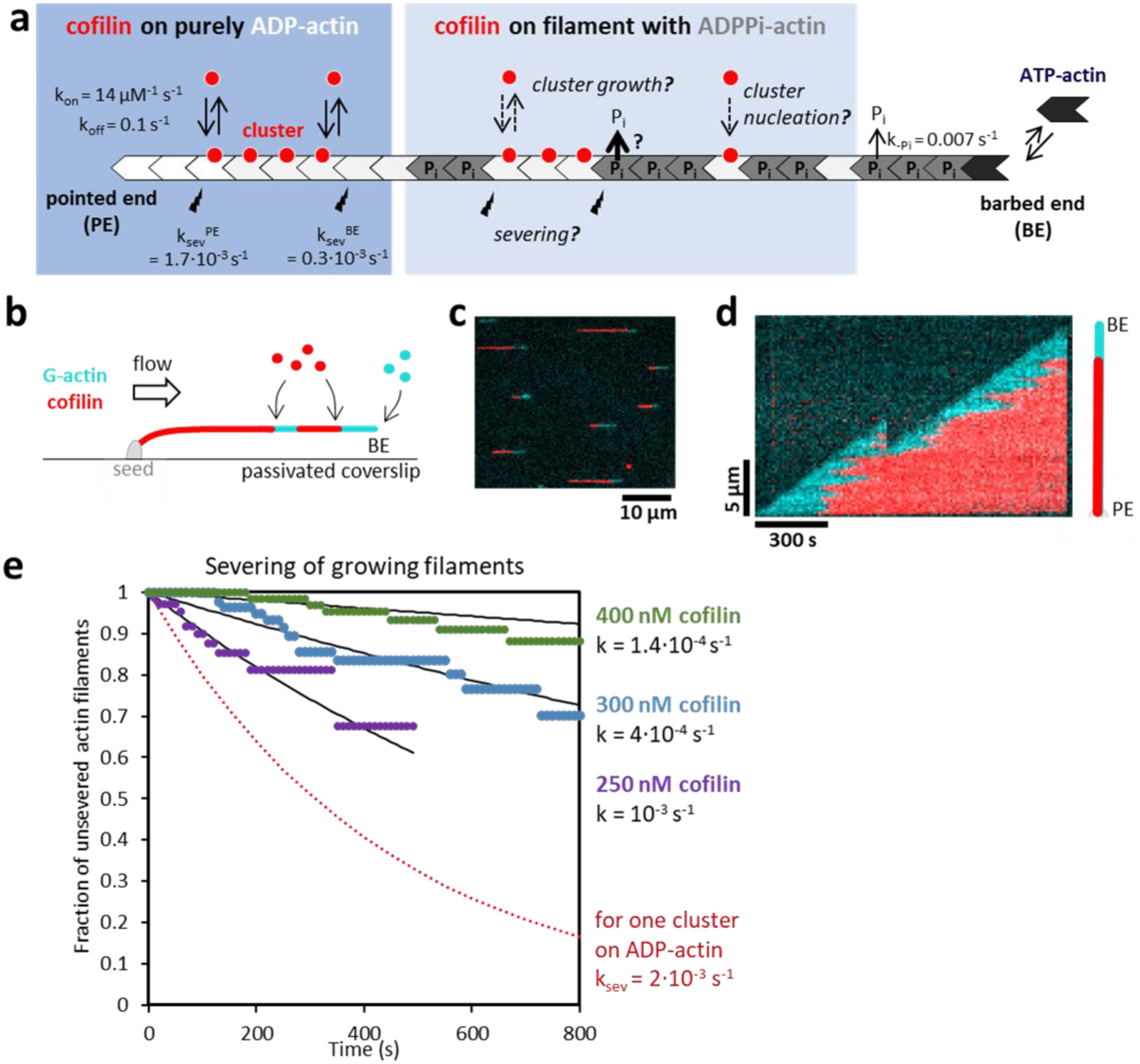
Actin filaments elongating while exposed to cofilin have a cofilin-free barbed end region, and sever slowly. **a.** Schematic description of cofilin binding to a growing actin filament. The rates for cofilin cluster growth and severing on ADP-actin are known (left), but the molecular events at play on filament regions containing ADPꞏP_i_-actin remain largely unexplored (center, light blue background). **b.** Sketch of a microfluidics experiment (side view). Filaments are elongated from surface-anchored spectrin-actin seeds. BE, barbed end. **c.** Typical fluorescence microscopy image of filaments exposed to 600 nM G-actin 10% Alexa 488 (cyan) and 400nM mCherry-cofilin1 (red) in a microfluidics chamber. Solutions flow from left to right. Only a fraction of the full field of view is shown. **d.** Kymograph of a filament from the experiment shown in b-c. Actin is in cyan and cofilin is in red. Images were taken every 10 seconds. A severing event can be seen (sudden shortening of the filament). **e.** Quantification of severing events over time, for growing actin filaments exposed to the indicated cofilin concentrations (250, 300 and 400 nM, n=30-60 filaments for each condition). The solid lines are exponential fits, yielding the indicated severing rates, per filament. For comparison, the dotted red line indicates the severing measured on ADP-actin, per cofilin cluster (Fig. S1b).

Our experiments were all done with purified proteins of mammalian origin (Methods). In particular, we used recombinant mouse mCherry-cofilin1 (cofilin, from hereon) and α-skeletal actin from rabbit muscle, 10% labeled on lysines with Alexa488 or ATTO488 (actin, from hereon). Experiments were performed at 25°C, at pH 7.4 with 50 mM KCl and 1 mM MgCl_2_. We made sure that flow rate and illumination, in the ranges that we used, did not impact the studied reactions.

As a reference, we first monitored the action of cofilin on ADP-actin. We grew filaments and aged them for 15 minutes (allowing more than 99.8% of subunits to become ADP-actin, given a P_i_-release rate of 0.007 s^−1^ (*4*, *5*)) before exposing them to cofilin. We found that cofilin clusters expanded symmetrically with rate constants k_on_= 14 µM^−1^s^−1^ and k_off_= 0.1 s^−1^ at cluster edges (Fig. S1a). We also monitored the severing of ADP-actin filaments, and found that severing occurred at a rate k_sev_= 2ꞏ10^−3^ s^−1^ per cluster (Fig S1b), and 84% of the time at the boundary growing toward the pointed end (k_sev_ ^PE^= 1.7ꞏ 10^−3^ s^−1^ ) and 16% at boundary growing toward the barbed end (k_sev_ ^BE^= 0.3ꞏ 10^−3^ s^−1^). These numbers are summarized on the sketch in Fig. 1a (left). They are consistent with our earlier measurements (*19*, *20*).

#### Growing actin filaments exposed to cofilin sever rarely

To investigate the action of cofilin on ADPꞏP_i_ -F-actin, we started by simply exposing growing actin filaments to cofilin (Fig. 1b-d). With high enough cofilin concentrations, these filaments become saturated by cofilin, except near their barbed ends. This has been observed before and previous reports referred to this bare region as the protective “ATP/ADPꞏP_i_ cap” (*32*).

Severing events occurred in the region partly decorated by cofilin, where cofilin cluster boundaries can be found, between the fully cofilin-saturated region and the cofilin-free region (Fig. 1d). We quantified the occurrence of severing events on these growing filaments and found that they were rare: filaments severed at rates ranging from 1.4ꞏ10^−4^ to 10^−3^ s^−1^ for cofilin concentrations ranging from 250 to 400 nM (Fig. 1e), while a single cofilin cluster on a fully ADP-actin filament severs at a rate of 2ꞏ10^−3^ s^−1^ (Fig. S1b, considering both boundaries). The number of cofilin cluster boundaries present on a growing actin filament is difficult to determine precisely, and it certainly depends on cofilin concentration, which could explain in part the cofilin-concentration dependence that we measured (Fig. 1e). However, we know there has to be at least one cofilin cluster boundary on growing actin filaments, and there seem to be several at the cofilin concentrations we used in our experiments (Fig. 1d). The severing rates we measured thus show that cofilin cluster boundaries on growing actin filaments sever more slowly than cluster boundaries on ADP-actin filaments.

#### Severing at ADPꞏP_i_-actin/cofilin cluster boundaries is very slow

Our result on the severing of growing actin filaments (Fig. 1e) suggests that an ADPꞏP_i_-actin/cofilin cluster boundary severs more slowly than an ADP-actin/cofilin cluster boundary. To test this hypothesis, we monitored the severing of individual cofilin clusters on actin filaments rich in ADPꞏP_i_ by first forming cofilin clusters on ADP-actin filaments and then exposing them to a solution containing inorganic phosphate (Fig. 2a-c). We found that the presence of P_i_ in solution reduced the rate of severing at cluster boundaries (Fig. 2c). This confirms that individual cluster boundaries sever more slowly when actin is in the ADPꞏP_i_ state. Notably, the presence of P_i_ did not seem to affect the asymmetry of severing, which remained far more frequent at the cluster boundary located toward the pointed end of the filament, as already observed on ADP-actin filaments (*19*, *25*, *26*).

**FIGURE 2.**
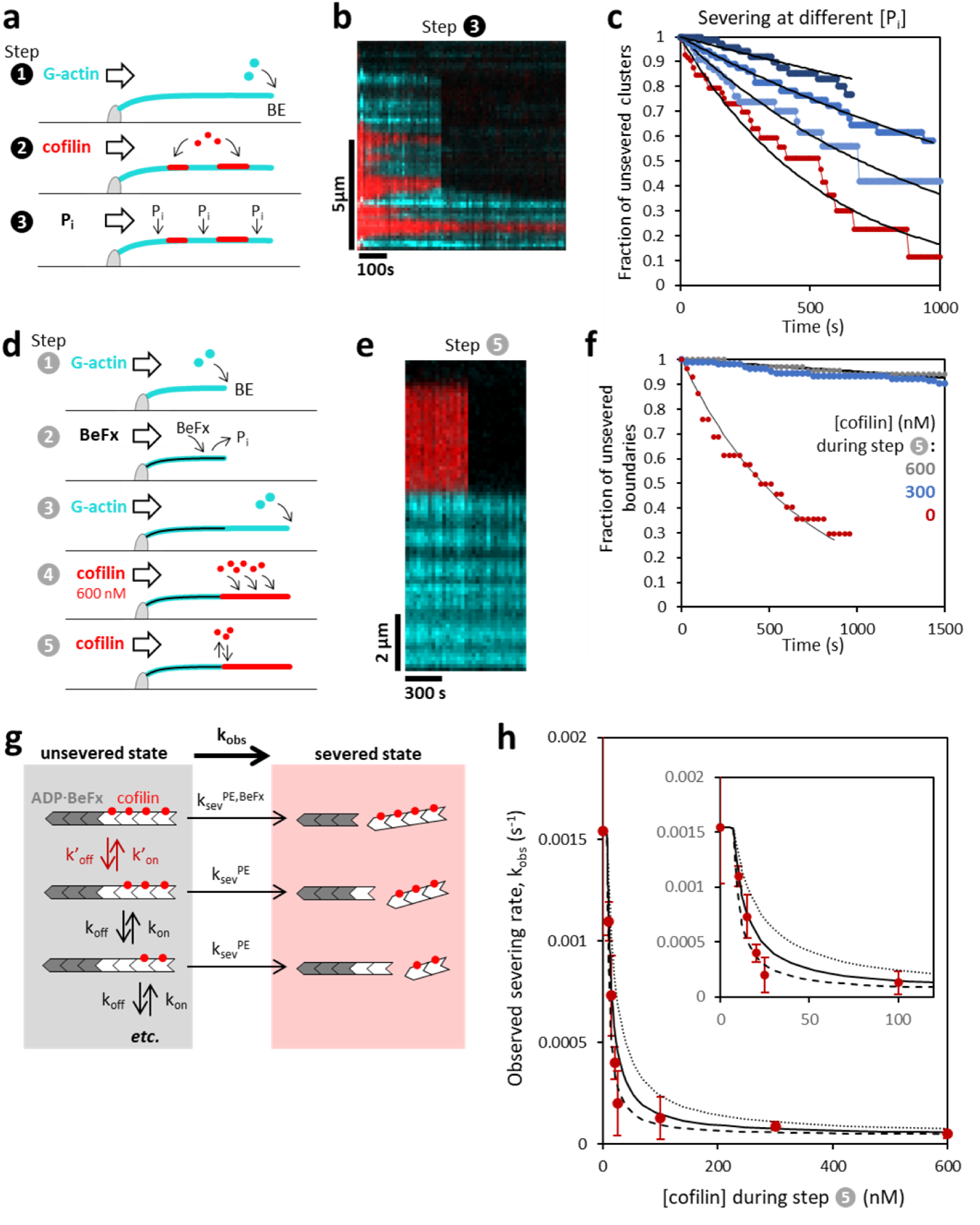
Severing and cofilin binding at ADPꞏP_i_-actin boundaries. **a-c. Severing at cofilin cluster boundaries in the presence of P_i_ in solution.** **a.** Sketch of the experimental sequence for the monitoring of severing events in the presence of P_i_. Filaments are elongated (step 1), then exposed to cofilin to form clusters (step 2), and finally exposed to buffer supplemented with Pi (step 3). Severing events, per cluster, are monitored during step 3. **b.** Kymograph of a single filament during step 3, exposed to 50 mM P_i_. Actin is in cyan and cofilin is in red. A severing event can be seen at a cluster boundary. **c.** Fraction of unsevered clusters (counting both boundaries) over time, while exposed to 0 (red), 5 mM (light blue), 10 mM (medium blue) and 25 mM P_i_ (dark blue), with exponential fits (black lines) yielding severing rates per cluster of 1.8 10^−3^, 10^−3^, 5.7 10^−4^, and 2.8 10^−4^ s^−1^, respectively. n=82-140 clusters in each condition. d-h. Severing at ADP-BeFx-actin/cofilin cluster boundaries. **d.** Sketch of the experimental sequence for the monitoring of severing events at BEFx-actin/cofilin cluster boundaries. Filaments are elongated (step 1) and converted to ADP-BeFx-actin (Step 2). A second filament segment is elongated (step 3) and saturated with cofilin (step 4). Severing is then monitored as the filaments are exposed to various concentrations of cofilin (step 5). **e.** Kymograph of a single filament during step 5, exposed to 4 nM cofilin. Actin is in cyan and cofilin is in red. The cofilin-free region (in cyan) is the ADP-BeFx-actin segment. A severing event can be seen at the cluster boundary. **f.** Fraction of unsevered cluster boundaries during step 5 of the experiment sketched in d. Filaments were exposed to 0 (red data points, n=30 filaments), 300 nM (blue, n=119) or 600 nM cofilin (gray, n=69). The thin black line is a fit of the data at 0 cofilin, yielding a rate of 1.5ꞏ10^−3^ s^−1^. The thick black line is an exponential fit of the data at 600 nM, yielding k ^PE,BeFx^=5ꞏ10^−5^ s^−1^. **g.** Schematic representation of the model for severing at the cluster boundary as a function of cofilin concentration. Cofilin (red dots) can only bind to ADP-actin subunits (white), with unknown rate constants k’_on_ and k’_off_ for the last ADP subunit (adjacent to the ADP-BeFx segment, in gray), and with known rate constants k_on_ and k_off_ for the other ADP subunits. When the last ADP subunit is occupied by cofilin, the cluster boundary severs at known rate k ^ADP,BEFx^. When it is unoccupied, the cluster boundary severs at known rate k ^PE^. **h.** Severing rate at the cluster boundary as a function of the cofilin concentration during step 5 of the experiment sketched in d. Each point is the average of n=2-4 experiments, except at [cofilin]=0 where n=24. Bars are standard deviations. Lines are computed based on the model sketched in g, with k’_on_=k_on_ and with k’_off_=k_off_ (thus K’_d_=k’_off_/k’_on_=7.1 nM, solid line), with k’_off_=2 k_off_ (K’_d_=14.3 nM, dotted line) and with k’_off_=0.5 k_off_ (K’_d_=3.6 nM, dashed line). Inset: same data with a different scale for cofilin concentration.

Since P_i_ can travel in and out of the nucleotide pocket of actin subunits, the severing rates we measured result from a combination of severing at ADPꞏPi- and ADP-actin boundaries. To quantify the severing rate at an ADPꞏP_i_-actin/cofilin cluster boundary, we used the phosphate analog beryllium fluoride (BeF_x_), which can replace P_i_ in the nucleotide pocket of actin subunits, from which it is released extremely slowly (*28*, *34*). ADPꞏBeF_x_-actin subunits in filaments thus mimic ADPꞏP_i_-actin which cannot release P_i_ and remain in the ADPꞏP_i_ state. We first formed fully ADPꞏBeF_x_-actin filaments, from which we then elongated a second segment by flowing ATP-G-actin (Fig. 2d). The second half of the filaments, devoid of BeF_x_, was then saturated with cofilin, and severing events at the ADPꞏBeF_x_-actin/cofilin cluster boundary were monitored (Fig. 2d-e). Note that we could not detect any cofilin on the ADPꞏBeF_x_-actin segment, and that severing events occuring elsewhere than at the cofilin cluster boundary were negligible. Consistent with our measurements in the presence of P_i_ in solution (Fig. 2a-c), we found a very low severing rate k ^PE,BeFx^ = 5ꞏ10^−5^ s^−1^ when exposing the filaments to high cofilin concentrations (Fig. 2f). This rate is ∼ 30-fold lower than the severing rate at ADP-actin/cofilin cluster boundaries on the PE side (Fig. S2).

### Cofilin has a high affinity for the binding site that borders a cluster and an ADPꞏBeF_x_-actin subunit

Our experiments on hybrid filaments, with one segment composed of ADPꞏBeF_x_-actin and another composed of ADP-actin, also allowed us to get insights into the affinity of cofilin for the ADP-actin subunit that neighbors the ADPꞏBeF_x_-actin region. We refer to this ADP-actin subunit as the last ADP-actin site. To get these insights, we monitored the severing of these hybrid filaments exposed to different concentrations of cofilin (Fig. 2g,h).

At high concentrations of cofilin, the last ADP-actin site is saturated, and severing occurs at the ADP- BeFx-actin/cofilin cluster boundary at the very slow rate k ^PE,BeFx^ (as in Fig. 2f). Switching to a solution that contains no cofilin, the last ADP-actin site becomes unoccupied, and severing occurs at an ADP-actin/cofilin cluster boundary, at a higher rate, matching the severing rate k_sev_ ^PE^ we measured for cofilin clusters on ADP-actin filaments (see sketch on Fig. 2g). The transition between these two extremes, at intermediate cofilin concentrations, allows us to estimate the affinity of cofilin for the last ADP-actin site that neighbors the ADP-BeFx-actin region. We find that this transition occurs at low cofilin concentrations, below 20 nM (Fig. 2h), which indicates that cofilin easily binds to the last ADP-actin site.

Comparing these data to a model assuming a rapid equilibrium for cofilin binding on the last ADP-actin site (Fig. 2g, Methods), we find it to be compatible with a dissociation constant in the range of a few nM (Fig. 2h). The affinity of cofilin for the site neighboring ADPꞏBeF_x_-actin is thus comparable to the affinity for an ADP-actin that does not neighbor ADPꞏBeF_x_-actin (K_d_=k_off_/k_on_=7.1 nM, based on the growth of a cofilin cluster on ADP-actin (Fig. S1a)). This result indicates that the binding of cofilin to an ADP-actin subunit is not affected by the presence of P_i_ on a neighboring subunit.

#### The nucleation of a cofilin cluster requires a minimal ADP-actin island

We next sought to investigate how cofilin clusters are nucleated on ADPꞏPi-rich actin filaments. However, since there is no established model for the simpler situation where cofilin clusters nucleate on purely ADP-actin filaments, we began by investigating this situation. We grew actin filaments, aged them for 15 minutes to let them become fully ADP-actin filaments, and exposed them to cofilin (Fig. 3a-c). We quantified the rate of nucleation of cofilin clusters per actin filament segment of 10 pixels, which corresponds to 481 actin subunits. We observed a non-linear increase of this nucleation rate k ^ADP^ as a function of cofilin concentration, over a 50-500 nM range (Fig. 3c).

**FIGURE 3.**
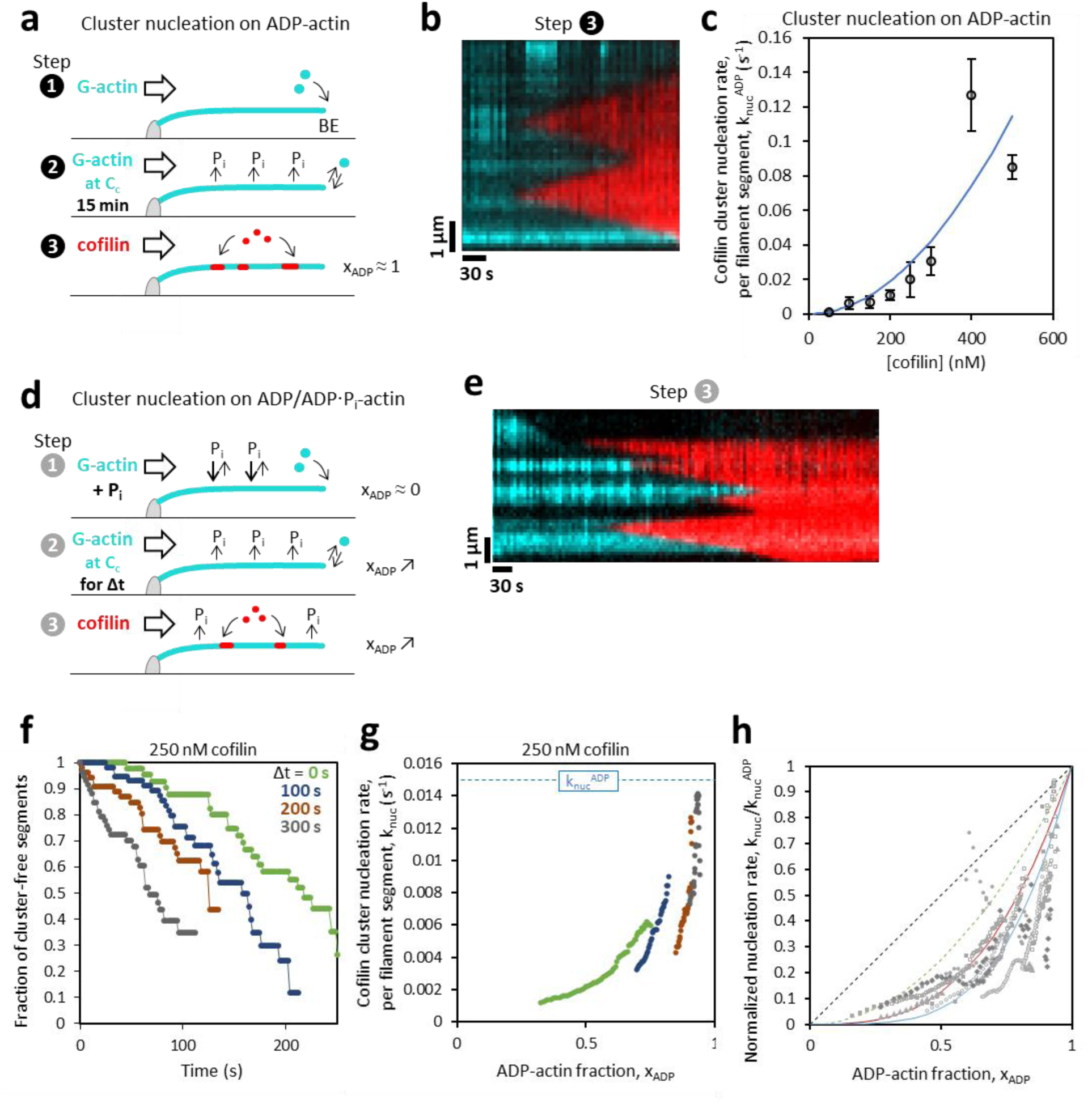
Nucleation of cofilin clusters on ADP-actin filaments and on ADP/ADPꞏP_i_-actin filaments. a-c. Cofilin cluster nucleation on ADP-actin filaments. **a.** Sketch of the experimental sequence for the monitoring of nucleation events on ADP-actin filaments. Filaments are elongated (step 1), then exposed to P_i_-free buffer (with G-actin at the barbed end critical concentration, C_c_) for 15 min to convert them entirely to ADP-actin (step 2), and finally exposed to various concentrations of cofilin (step 3). Nucleation events are monitored during step 3. **b.** Kymograph of a single filament during step 3, exposed to 100 nM cofilin. Actin is in cyan and cofilin is in red. Two nucleation events are visible. **c.** Cofilin cluster nucleation rate, k_nuc_^ADP^, per ADP-actin filament segment of 1.3 µm, measured for different cofilin concentrations (step 3 of the experiment sketched in a). Each point is the average of 2-7 independent measurements, each on a minimum of 30 filament segments. Bars are standard deviations. The blue line is a theoretical curve computed with k_on_^single^ = 0.03 µM^−1^ s^−1^ and k_off_^single^ = 0.9 s^−1^ (Methods). d-h. Cofilin cluster nucleation on ADP/ADPꞏP_i_-actin filaments. **d.** Sketch of the experimental sequence for the monitoring of nucleation events on filaments with different ADP-actin fractions x_ADP_. The filaments are elongated in the presence of 80 mM P_i_ to maintain them in the ADPꞏP_i_ state (step 1), then exposed to P_i_-free buffer (with G-actin at the barbed end critical concentration, C_c_) for a time Δt to partly convert them to ADP-actin (step 2, x_ADP_ increases), and finally exposed to various concentrations of cofilin (step 3, x_ADP_ continues to increase). Nucleation events are monitored during step 3. **e.** Kymograph of a single filament during step 3. Actin is in cyan and cofilin is in red. Several nucleation events are visible. **f.** Survival functions for the nucleation of clusters on 1.3-µm filament segments (n=50-60 filament segments), exposed to 250 nM cofilin during step 3, after different delays Δt during step 2. **g.** Cofilin cluster nucleation rates k_nuc_ per 1.3-µm filament segment, with ADP-actin fraction x_ADP_, derived from the survival functions shown in f (Methods). The dashed line indicates the nucleation rate measured on ADP-actin filaments with the same cofilin concentration. **h.** Normalized nucleation rate, k_nuc_/k ^ADP^, as a function of the ADP-actin fraction x_ADP_, from 6 independent experiments at different cofilin concentrations: 150 nM (gray dots, triangles), 200 nM (open circles, gray squares, diamonds), and 250 nM cofilin (open squares, same data as in g). The lines are power functions y=x^p^, with p=1 (black, dashed), p=2 (green, dashed), p=3 (red) and p=4 (blue).

Based on cryo-EM data, Huehn et al. (*22*) proposed that cooperativity in cofilin binding to actin fiaments is effective when at least two cofilins are bound on neighboring sites. We thus compared our data to a model assuming that the first cofilin binds with rate constants k_on_^single^ and k_off_^single^, and that the independent binding of a second cofilin, also with rate constant k_on_^single^, on the neighboring site on the other strand of the filament, will stabilize their interaction with the filament. The cofilin doublet thus formed behaves as a cluster, and expands with the rate constants k_on_ and k_off_ that we determined for larger cofilin clusters (Fig. S1a). Fitting our data with this model did not yield a unique pair of values for k_on_^single^ and k_off_^single^ because these two parameters are strongly correlated (Methods). To constrain these parameters, we turned to previous measurements by Hayakawa et al. who monitored the binding of single cofilins using a slightly different fluorescent cofilin construct at pH 7.0, and found on- an off-rate constants of 0.06 µM^−1^s^−1^ and 0.6 s^−1^, respectively (*24*). Our data can be well described by the computed curve using values that are remarkably close : k_on_^single^ = 0.03 µM^−1^ s^−1^ and k_off_^single^ = 0.9 s^−1^. (Fig. 3c, Methods).

Our data is thus consistent with a model for cofilin cluster nucleation on ADP-actin where 2 cofilins have to find themselves, by chance, on two contiguous binding sites on the two strands of the filament. Once this cofilin doublet is formed, it behaves like a cluster, with a smaller off-rate (0.1 versus 0.9 s^−1^) and a much higher on-rate constant for the addition of more cofilins (14 versus 0.03 µM^−1^s^−1^) compared to isolated cofilins.

We next monitored the nucleation of cofilin clusters on filaments containing ADPꞏP_i_-actin (Fig. 3d-e). To do so, actin filaments were exposed to a solution of 80 mM Pi for a few minutes, to make them nearly fully ADPꞏP_i_-actin. They were then exposed to regular, P_i_-free buffer for a duration Δt, ranging from 0 to 400 seconds. During that time, actin subunits began releasing their Pi at rate k_-Pi_=0.007 s^−1^ corresponding to a half-life of ∼100 seconds (*4*, *5*). After that time, we exposed the filaments to cofilin, also in P_i_-free buffer. This allowed us to monitor the nucleation of cofilin clusters on filaments that had different average contents in ADPꞏP_i_-actin, which we quantified with the ADP-actin fraction, x_ADP_=1- exp(-k_-Pi_ꞏt), where t is the time since the filaments were switched to P_i_-free buffer.

As expected, cofilin clusters nucleated faster on filaments that were allowed to release their P_i_ for a longer time Δt before being exposed to cofilin (Fig. 3f). These survival curves are not exponential because the rate of nucleation is not constant but evolves over time, as more P_i_ is being released from the filaments. We estimated the nucleation rate k_nuc_(t) at different time points, corresponding to different values of x_ADP_ (Methods) and found that k_nuc_(x_ADP_) had a concave shape, first increasing slowly then sharply as x_ADP_ increases (Fig. 3g).

We repeated this experiment for different cofilin concentrations and found that the nucleation rate k_nuc_ always showed a similar increase as a function of x_ADP_. This can be evidenced by plotting, for each cofilin concentration, the nucleation rate k_nuc_ normalized by its maximum value k_nuc_^ADP^ (determined by an exponential fit of the survival curve for nucleation on filaments for which Δt=900s, and thus x_ADP_>0.998) as a function of x_ADP_ (Fig. 3h). We compared these normalized nucleation rates to power laws y= x_ADP_^p^, which represent the fraction of actin subunits that are in the ADP state and are surrounded by (p-1) ADP-actin subunits. We found that the normalized nucleation rates are globally close to the curves computed with p=3 and p=4 (Fig. 3h).

This means that the rate at which clusters nucleate is equal to the nucleation rate on ADP-actin times the density of ADP-actin subunits that are part of “ADP islands”, comprising at least 3-4 contiguous ADP-actin subunits.

This is consistent with the notion that the limiting step for cluster nucleation is the formation of a minimal cluster of two cofilins. It is also consistent with a model where the only thing reducing the nucleation of cofilin clusters on ADPꞏP_i_-containing actin filaments is the availability of large enough islands of contiguous ADP-actin, that form spontaneously as actin releases its P_i_. On these islands, nucleation thus seems unhindered by the nearby ADPꞏP_i_-actin subunits, which is reminiscent of our finding that cofilin binding at cluster boundaries is unaffected by the presence of BeF_x_ on the next subunit (Fig. 2g-h).

#### Cofilin clusters grow slower and asymmetrically in the presence of ADPꞏP_i_-actin subunits

Using a strategy similar to the one used to study cluster nucleation, we monitored the growth of cofilin clusters on filaments with different ADPꞏP_i_/ADP contents (Methods). We find that in the presence of ADPꞏP_i_-actin, clusters grow more slowly and asymmetrically, with a faster expansion toward the pointed end (Fig. 4a). We did not observe this asymmetry when the filaments were entirely made of ADP-actin (Fig. 4b), consistent with our previous observations on ADP-actin filaments (*19*).

**FIGURE 4.**
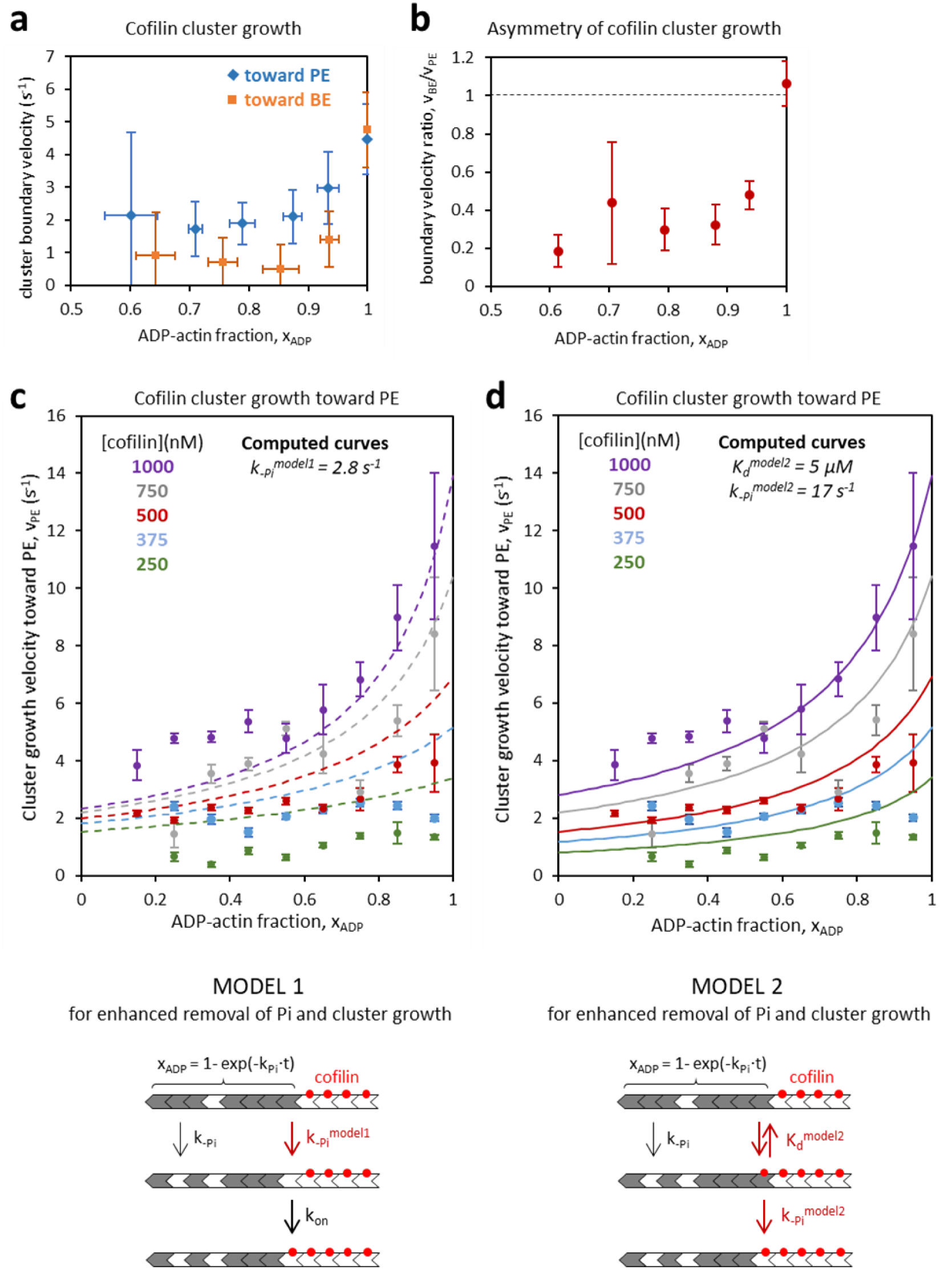
Expansion of cofilin clusters on filaments containing ADP- and ADPꞏP_i_-actin. **a.** Velocity of cluster growth, for boundaries in both directions, as a function of the ADP-actin fraction x_ADP_. Data points from 5 independent experiments were pooled according to their values of x_ADP_, and averaged (for each value of x_ADP_, n=6-20 boundaries toward the PE, and n=9-23 boundaries toward the BE). Bars are standard deviations. **b.** Asymmetry of cluster growth, expressed as the ratio of velocities v_BE_/v_PE_, as a function of ADP-actin fraction x_ADP_ (based on the data from a). Bars are standard errors. The dashed line indicates symmetrical cluster expansion, with a velocity ratio of 1. **c.** Velocity of cluster growth toward the PE, as a function of the ADP-actin fraction x_ADP_, for different cofilin concentrations. Dashed lines are computed curves based on “model 1”, summarized on the schematic below, for the different cofilin concentrations. In model 1, the only free parameter is the rate k_-Pi_^model1^, at which P_i_ is released from an ADPꞏP_i_-actin at a cofilin cluster boundary (Methods). **d.** Same experimental data as in c. The solid lines are computed curves based on “model 2”, summarized on the schematic below, for the different cofilin concentrations. In model 2, there are two free parameters: the dissociation constant for cofilin binding to ADPꞏP_i_-actin at a cluster boundary, K_d_^model2^, and the rate of P_i_ release from the cofilin-bound ADPꞏP_i_-actin, k_-Pi_^model2^ (Methods). See also Fig. 5b.

To further analyze the impact of ADPꞏP_i_-actin on cluster growth, we focused on the faster boundary, the one moving toward the pointed end. We measured the cluster growth velocity at this boundary, v_PE_, for different ADP-actin fractions x_ADP_, and for different cofilin concentrations (Fig. 4c).

We first tried to fit this set of data with the classical model (*32*), which we refer to as “model 1”, where the release of P_i_ from an ADPꞏP_i_-actin subunit is enhanced by the presence of cofilin on a neighboring ADP-actin subunit (sketch in Fig. 4c, Methods). The only free paramater is k ^model1^, the enhanced P_i_ release rate. We found that this model could not account well for our data: varying k ^model1^ allowed to tune the amplitude of the velocity change over the different values of x_ADP_, but it could not reproduce the differences in velocity that we measured at low values of x_ADP_ for different cofilin concentrations (Fig. 4c, Fig. S3). Importantly, modifying the model by increasing the range over which P_i_ release is enhanced, more than one subunit away from the nearest cofilin, would not allow to amplify differences between cofilin concentrations at low x_ADP_. On the contrary, increasing the range would favor the expansion of the slowly growing clusters, because ADPꞏP_i_-actin subunits would spend more time within the range, thereby increasing their chances to become ADP-actin before being reached by the cluster boundary.

We reasoned that the model was missing a cofilin-concentration dependent step that would be involved in the release of P_i_, in order to account for the differences between cofilin concentrations that we measured at low x_ADP_. We thus considered a different model, which we call “model 2”, where cofilin would be able to bind an ADPꞏP_i_-actin subunit in an unstable state, from which either cofilin or P_i_ is rapidly released (sketch in Fig. 4d, Fig. 5b, Methods). The possibility to have both cofilin and P_i_ bound to the same actin subunit is consistent with our observation that P_i_ in solution accelerates the rate of cofilin cluster disassembly (Fig. S2). Indeed, since the affinity of cofilin for ADP-actin at a cluster boundary is not affected by the presence of P_i_ in the neighboring subunit (Fig. 2g-h), it is unlikely that P_i_ in solution would accelerate the dissociation of cofilin by binding to its ADP-actin neighbor. Instead, a likely explanation would be that P_i_ is able to enter the nucleotide pocket of the cofilin-bound ADP-actin subunit at the cluster boundary, thereby accelerating the dissociation of cofilin.

**FIGURE 5.**
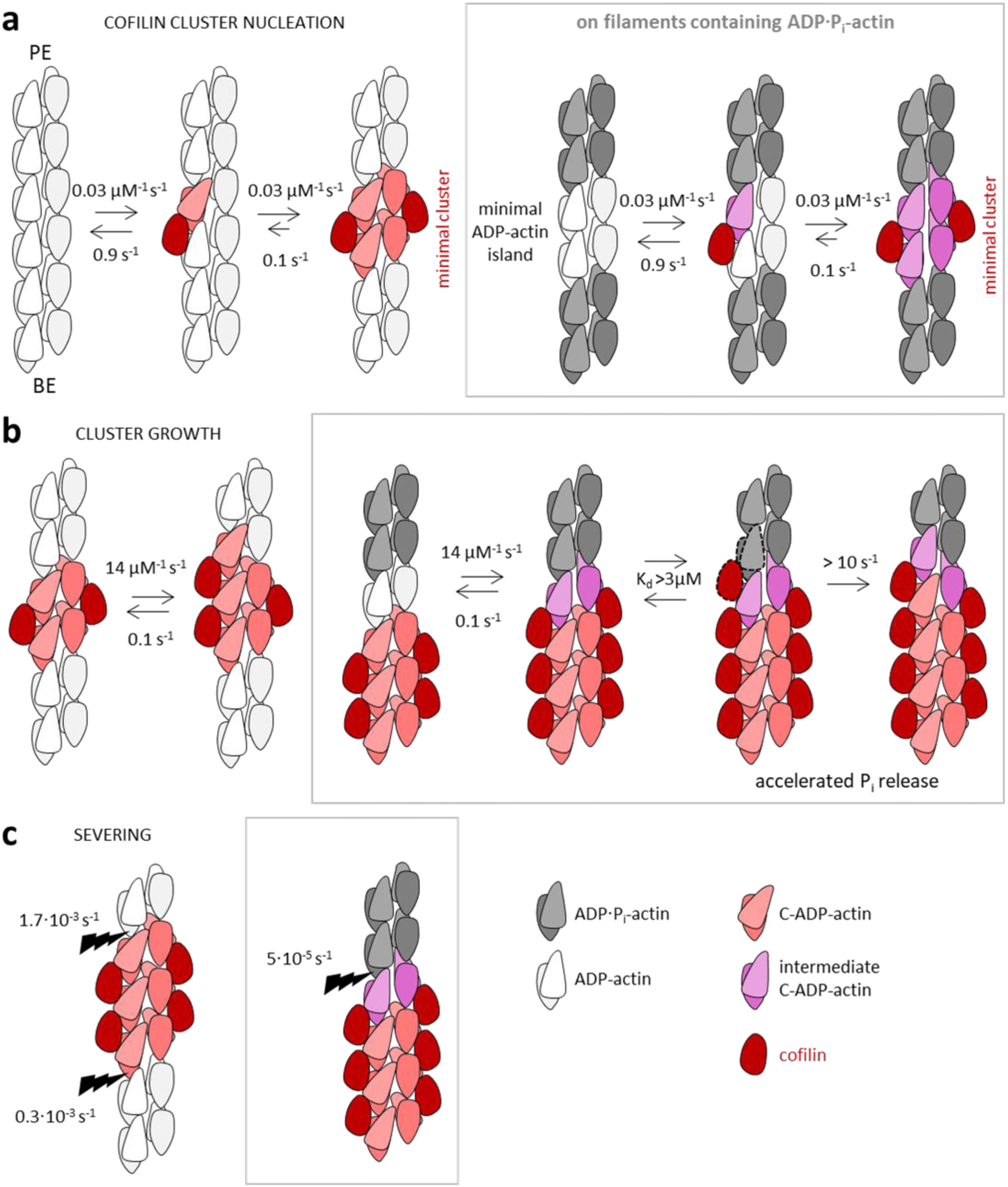
Summary and interpretation of results. Shematics recapitulating the rates and mechanisms from this study, for cofilin cluser nucleation. (a) and growth (b), as well as their ability to sever actin filaments (c), as a function of their nucleotide content. The structural interpretation and representation are based on Huehn et al. (*22*). The two strands of the actin filament are represented flat, without the helical twist. For each actin subunit, the two domains are schematized, the lighter-colored being the outer domain. Cofilin binds two actin subunits, and it cannot bind to ADPꞏP_i_-actin (in gray) but only to ADP-actin (in white). The binding of cofilin causes a conformational change in actin, consisting mostly in a tilt of the outer domain of the subunit (C-ADP-actin conformation, in red). This weakens the bonds with longitudinal neighbors, especially in the PE direction. The binding of a single cofilin changes the conformation of only one actin subunit (a) (*22*).Two bound cofilins form a minimal cluster. We propose that ADP-actin subunits bound simulateously to cofilin and to an ADPꞏP_i_-actin neighbor adopt an intermediate conformation (shown in purple), where the outer domain is not as tilted as in the C-ADP-actin conformation. In this intermediate conformation, the affinity for cofilin remains high (binding rates are comparable to standard ADP-actin, as shown in a and b), while the interaction with the longitudinal neighbor is less affected than with the C-ADP-actin conformation (explaining why this interface severs slowly, as shown in c). When a cluster grows, we propose that cofilin is able to interact weakly (high dissociation constant) with ADPꞏP_i_-actin at the cluster boundary (dashed outlines in b). This unstable state promotes the rapid release of P_i_ from the ADPꞏP_i_-actin.

We thus tried to fit our data with model 2, assuming that cofilin is in rapid equilibrium with the ADPPi- actin subunit at the cluster boundary, with a dissociation constant K_D_^model2^, and with a rate k ^model2^ for the release of P_i_ from the cofilin-bound ADPꞏP_i_-actin subunit (Fig. 4d, Methods). K_D_^model2^ and k ^model2^ are the only free parameters, and they appear to be strongly correlated in the range that fits our data well. We find that the model is in best agreement with our data for a ratio K_D_^model2^ / k ^model2^ ∼0.3 µMꞏs and values larger than K_D_^model2^ = 3 µM (and k ^model2^ =10 s^−1^). Larger values, keeping the ratio constant, do not improve the fit much (Fig. S4). Fig. 4d shows the curves computed with K ^model2^=5 µM (k ^model2^ =17 s^−1^).

In summary, our data for the acceleration of cofilin cluster growth toward the PE as the filament ages is compatible with a model where cofilin binds to ADPꞏP_i_-actin at the cluster boundary with a low affinity (dissociation constant in the µM range or above), and where P_i_ is rapidly released from the cofilin-bound ADPꞏP_i_-actin subunit (orders of magnitude faster than in the absence of cofilin).

## Discussion

By creating specific, controlled conditions for single actin filaments, we have isolated individual reactions, allowing us to obtain mechanistic insights and to quantify reaction rates for the interaction of cofilin and actin filaments.

While our objective was to focus on filaments containing ADPꞏP_i_-actin, our results also shed light on some of the questions that remained unanswered regarding the action of cofilin on ADP-actin filaments. For instance, our kinetic study of nucleation agrees with a model proposed by Huehn et al. (*22*) based on cryo-EM observations, where two cofilins need to bind on two contiguous sites on the two strands of the actin filament to form a minimal cluster and initiate the cooperative binding of more cofilins on the adjacent sites (Fig. 5a). In contrast, our results on filament severing at ADPPi- actin/cofilin cluster boundaries (Fig. 2) contradict the mechanical model for severing. Indeed, this model proposes that severing is primarily driven by the mechanical difference between the softer cofilin-decorated segment and the stiffer bare ADP-actin segment. It would thus predict a faster severing when the latter is replaced by an even stiffer ADP-BeFx-actin segment (*35*). We find the opposite. Finally, our observation that the presence of Pi in solution and in the actin subunits causes cofilin clusters to grow faster toward the pointed end (Fig. 4a,b) may provide clues explaining why some experimental configurations also lead to an asymmetric growth of cofilin clusters on ADP-actin filaments.

Our results on filaments that contain ADPꞏP_i_ -actin have led us to propose mechanisms for the nucleation and the growth of cofilin clusters on these filaments. They explain why cofilin is unable to catch up with the growing BE of an actin filament (*32*), as cluster nucleation is limited by the spontaneous appearance of ADP-actin islands (Fig. 3h), and clusters then grow slowly toward the BE (Fig 4a,b). Importantly, the picture that emerges from our different results is one where the impact of the nucleotide state remains very local: cofilin clusters nucleate on ADP islands without being affected by the surrounding ADPꞏP_i_-actin subunits (Fig. 3h) ; at cluster boundaries, the affinity of cofilin for the last ADP-actin at the edge of a segment of ADPꞏBeFx-actin is barely affected by the presence of this segment (Fig. 2h) ; and clusters grow by enhancing Pi release from the nearest ADPꞏP_i_-actin only (Fig. 4c,d). Together, our results thus argue in favor of nearest-neighbor cooperativity, in agreement with indications from cryo-EM data (*16*, *22*).

It is important to note that the models we have used to fit our data are all very simple, with a minimal set of free parameters. Our objective was to test whether the main trends of the data could be captured by making simple assumptions. For the growth of cofilin clusters, for example, our proposal is that cofilin-ADPꞏP_i_-actin likely exists as an unstable state (model 2, Fig 4d) and that this could be key, but it does not exclude the possibility for cofilin to also enhance Pi release from an neighboring ADPPi- actin without binding to it (as in model 1, Fig 4c). Similarly, our model for nucleation (Fig 3c) considers that a cofilin doublet behaves like larger clusters, with the same on- and off-rates at their boundaries, but it may be that cofilin doublets are a bit less stable than larger clusters.

For these models, we have used a one-strand depiction of the actin filaments, and did not specify that cofilin binds two actin subunits. Nonetheless, based on the available structural data for cofilin on actin filaments, we can propose a higher-resolution interpretation of our results (Fig. 5). When cofilin binds to the side of an ADP-actin filament, the two subunits in contact with cofilin adopt a more twisted conformation, mainly due to the tilting of actin’s outer domain, which weakens its bonds with its neighboring actin subunits (*15*, *22*). Our data suggest that an ADP-actin subunit in contact both with cofilin and with an ADPꞏP_i_-actin neighbor adopts a different conformation, where the bonds with the ADPꞏP_i_-actin neighbor are mostly maintained with almost no penalty for the binding of cofilin (Fig 5b,c). Such an “intermediate” conformation would explain the low severing rate we measured for ADPꞏP_i_- actin/cofilin cluster boundaries (Fig. 2) as well as the high affinity of cofilin for ADP-actins that neighbor an ADPꞏBeFx-actin (Fig. 2,h). In contrast, in the unstable state where cofilin would bind to an ADPꞏP_i_- actin subunit, that we propose to explain our cluster growth data (Fig 4c,d), the conformation of the actin subunit likely remains unfavorable to cofilin binding (Fig. 5b). Future cryo-EM studies, as well as molecular dynamics studies, investigating the structure of ADPꞏP_i_-actin/cofilin cluster boundaries, should be able to test the existence of the new conformations we propose.

In cells, filaments containing ADPꞏP_i_-actin are a relevant substrate to consider for actin-binding proteins, because P_i_ release takes some time after a filament has assembled (unless accelerated by proteins or other factors), and because P_i_ is expected to be in the mM range in the cytosol of most cell types. This makes it unlikely for filaments to become fully ADP-actin in cells without the intervention of regulatory proteins. In fact, future studies should consider situations where Pi is present in solution and can bind to ADP-actin subunits, in competition with cofilin.

Importantly, several actin-binding proteins are present in cells besides cofilin, and some are also able to sense and modify the nucleotide state of actin. Coronin, for example, binds to ADPꞏP_i_-actin filaments, favors the release of Pi and the recruitment of cofilin (*36*, *37*). This mechanism seems to provide another means for cofilin to rapidly decorate filaments that are initially rich in ADPꞏP_i_-actin. We hope our results will serve as a base, on which future studies will build to investigate the regulation of actin filament disassembly in the full complexity of their nucleotide state.

## Acknowledgements

The literature on cofilin is vast and we apologize to the authors whose work we could not cite. We thank Ingrid Billault-Chaumartin and LuYan Cao for their constructive reading of this manuscript.

## Funding

We acknowledge funding from the Agence Nationale de la Recherche (grant ANR-21-CE13- 0043-01 to G.R.-L.), from the Fondation pour la Recherche Médicale (grant EQU202203014630 to G.R.- L.) and from the Fondation Bettencourt-Schueller (grant Impulscience 1235 to A.J.).

## METHODS

### Proteins

Actin. α-skeletal muscle actin (Uniprot P68135) was purified from rabbit muscle acetone powder by standard polymerization/depolymerization cycles (*38*). Actin was labelled on surface lysines with Alexa-488 or ATTO-488-sulfoNHS.

Cofilin. Mouse cofilin-1 (Uniprot P18760) was expressed in E. coli as a GST fusion, cleaved with PreScission protease, and purified by gel filtration (*39*). Cofilin was N-terminally tagged with mCherry via a 6-amino-acid linker.

Profilin. Human profilin-1 (Uniprot P07737) was purified by poly-L-proline affinity chromatography (*40*) and used in all filament polymerization steps to prevent spontaneous nucleation of filaments in solution.

Spectrin-actin seeds, used to anchor filaments to the coverslip from their pointed ends and allow their elongation from the barbed end, were purified from human erythrocytes (*41*). More details on these protocols can be found in (*19*).

### Buffers

Most steps were done in standard F-buffer (5 mM Tris-HCl, 1 mM MgCl₂, 0.2 mM EGTA, 0.2 mM ATP, 10 mM DTT, 1 mM DABCO, 0.1% BSA, 50 mM KCl, pH 7.4).

Steps with BeFₓ (10 mM NaF + 2 mM BeSO₄) used the same buffer adjusted to pH 6.6.

A Pi buffer, prepared by replacing KCl with 30.75 mM K₂HPO₄ and 19.25 mM KH₂PO₄ (50 mM Pᵢ total, pH 7.4), was used to expose filaments to inorganic phosphate. To keep the ionic strength constant, the Pi concentration was varied below 50 mM, and the KCl concentration was adjusted accordingly. To saturate filaments with P_i_, however, we used 80 mM P_i_ and the ionic strength was thus higher during this step.

### Experiments

#### Microfluidic chambers

were made from PDMS (10:1 PDMS:curing agent, cured 2 h at 70°C) using pre-existing wafer molds, yielding 20 µm-high, ∼1 cm-long chambers with three or four inlets converging into a single central channel and outlet. Chambers were bonded to mica glass coverslips, pre-cleaned by sonication in Hellmanex followed by successive treatment with 1 M KOH and ethanol, using UV-activated surface bonding. The chamber inlets were connected to tubes in which pressure was controlled using an MFCS-EZ device (Fluigent), and the resulting flow rates were measured using Flow units (Fluigent).

#### Surface seeding and passivation

Chambers were seeded with ∼5 pM spectrin-actin seeds, adsorbed non-specifically to the glass coverslip for 1 min 30 s, then passivated with 50 mg/mL BSA for 15 min; for experiments requiring better passivation (i.e. involving BeFx or P_i_), an additional 15 min passivation with casein (1.25 mg/mL) and F127 (1 mg/mL) was done.

#### Microscope: wide-field or TIRF

Imaging was performed on Nikon TiE, Ti2 and TE2000 inverted microscopes with TIRF 1.49 NA 60× and 100x oil-immersion objectives, using TIRF, HiLo, or epifluorescence illumination depending on the experiment. The TiE and Ti2 systems were illuminated with tunable lasers (100 mW and 150 mW) and a TIRF set-up (iLAS2, Gataca Systems) controlled using Metamorph (TiE) or micromanager (Ti2), and images were acquired using an Evolve EMCCD camera (Photometrics)(TiE) or an sCMOS-kinetix camera (photometrics)(Ti2). The TE2000 system was controlled with Micro-Manager, illuminated with an Xcite lamp (120 W), and images acquired on an Orca-Flash 4.0 sCMOS camera (Hamamatsu). All microscopes used a a motorized stage (Marzhauser), and experiments were performed at 25°C.

#### Typical filament elongation experiment

Filaments were elongated from surface-anchored spectrin-actin seeds by flowing 0.6 to 1 µM G-actin (10% labelled) with equimolar or slightly higher concentrations of profilin (0.6 to 1.2 µM).

#### Reproducibility

Experimental conditions were controlled as much as possible to ensure reproducibility. Different batches of cofilin were prepared over the course of this project, and their behaviors were indistinguishable. However, some aliquots from the two latest batches exhibited cluster nucleation and growth rates that were 2-3 times faster than our standard, observed with the majority of aliquots. We could not find an explanation for their behavior. We treated them as outliers, and discarded the results obtained using these aliquots.

### Image analysis

Images were analyzed using Fiji (ImageJ). Movies were registered, to correct for the jitter sometimes induced by the multi-position acquisitions. To measure the cluster growth velocity, we manually tracked the position of the half-maximum intensity on the mCherry-cofilin1 fluorescence intensity profile, in Fiji. For the rapidly growing clusters (above 100 nM cofilin, on ADP-actin) we directly measured the slope (angle) on kymographs in Fiji. The nucleation of cofilin clusters was detected manually, either on kymographs or on the plot intensity profiles. Severing events were readily detected, on the movies or on the kymographs.

The observation of individual events (nucleation, or severing) were used to compute survival functions. Censoring events were taken into account using a Kaplan-Meier algorithm.

### Data analysis and modeling

We present here the models we compared to our experimental data. In one case (severing at ADPꞏBeFx-actin/cofilin cluster boundary, see below) we could not solve our model analytically and we used simulations. When we could derive an analytical solution, we used the curve_fit function from the SciPy package in Pyhton to compare the computed curves to our data. It also yielded the covariance matrix, which indicated how correlated our free parameters were.

#### Severing at ADPꞏBeFx-actin/cofilin cluster boundary (*Fig. 2d-h*)

Severing occurs at slow rate *k_sev_ ^BeFx^* when there is ADPꞏBeFx-actin at the cofilin cluster boundary, and at a faster rate *k_sev_^PE^* when there is ADP-actin at the cofilin cluster boundary (as sketched in Fig. 2g). The first situation is obtained with high [cofilin], while the second is obtained when removing cofilin from solution, causing the cluster to shrink and thus expose ADP-actin at its boundary (i.e. [cofilin]=0 in step 5 of the experiment, Fig. 2d). For intermediate [cofilin], the observed severing rate *k_obs_* will depend on the state of the cluster boundary, that evolves over time as cofilin is added and removed. We assumed that the two states for the cluster boundary were in rapid equililbrium, and computed *k_obs_* as the sum of the two rates *k_sev_^PE,BeFx^* and *k_sev_ ^PE^* weighted by the fraction of time spent in each state.

We ran a simple Monte-Carlo simulation, using a Gillespie algorithm, to compute how long the cluster boundary is at an ADP-actin subunit (i.e. how long it takes for the boundary to come back to the state where it neighbors ADPꞏBeFx-actin subunit), for different cofilin concentrations. This computation depends on the on- and off-rate constants for cofilin binding to a regular ADP-actin subunit (k_on_ and k_off_), which have fixed values that we determined independently (Fig. S1a), and on the on- and off-rate constants for cofilin binding to the ADP-actin subunit that neighbors ADPꞏBeFx-actin (k’_on_ and k’_off_), which are unknown.

#### Cluster nucleation on ADP-actin filaments (Fig. 3a-c )

We compute the rate of formation of cofilin doublets, based on the reaction scheme sketched on Fig. 5a, as follows. We consider that individual cofilin molecules are in rapid equilibrium between being free in solution and bound as a single cofilin on the actin filament, with rate constants *k_on_ ^single^*and *k_off_^single^* (and a dissociation constant *K_d_^single^= k_off_^single^ / k_on_ ^single^* ). We consider filament segments of N actin subunts, corresponding to N binding sites for cofilin (in the analysis of our experimental data, we chose segments of 1.3 µm, so N=481 actin subunits).

On average, there are thus *Nꞏ[cofilin]/( K_d_^single^ + [cofilin])* individual cofilins on the segment. Each of these cofilin singlets can become a doulet if another cofilin binds to the neighboring site on the other strand of the actin filament. The rate at which cofilin doublets are formed on the filament segment can thus be written as *Nꞏk ^single^[cofilin]^2^/( K_d_ ^single^ + [cofilin])*.

The cofilin doublet is assumed to behave as a cluster, adding or losing cofilins at its boundaries with known rate constants *k_on_* and *k_off_*. Its length fluctuates and there is a probability for a small cluster to disassemble before it can be observed in our experiments. The probablity for a cofilin cluster to lose a cofilin before adding one is *p_shrink_=k_off_/(k_off_+k_on_[cofilin])*. With rate constants *k_on_* = 14 µM^−1^s^−1^ and *k_off_*=0.1 s^−1^ (Fig. S1a), this probability is very low in the range of cofilin concentrations we have used (for example, *p_shrink_*= 0.125 at 50 nM, 0.067 at 100 nM and 0.023 at 300 nM cofilin). We have computed that, in our experimental conditions, the probability for a cofilin doublet to reach a size we can observe is close to 1.

We thus estimate the cofilin cluster nucleation rate on ADP-actin segments of N subunits to be:

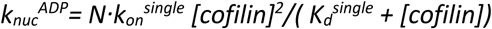

This is the equation used to compute the curve shown in Fig. 3c.

#### Cluster nucleation on ADPꞏP_i_-contaning actin filaments (Fig. 3f-h)

The survival curves (Fig. 3f) are not exponential because the rate of nucleation is not constant, but increases over time as the nucleotide content of the filaments evolve. The rate of nucleation at time t can be determined from the survival curve as *-S’(t)/S(t)* where *S(t)* is the survival function and *S’(t)* is its time derivative at time t, which we estimated by a local linear fit of the survival curve. The average ADP-actin fraction ADP-actin fraction of the filament at time t was computed as x_ADP_=1-exp(-k_-Pi_ꞏt), where t is the time since the filaments were switched to P_i_-free buffer (i.e. including the delay Δt during step 2 in Fig. 3d).

#### Cluster growth on ADPꞏP_i_-contaning actin filaments (Fig 4c,d)

We compute the average velocity of the PE-side cluster boundary as *v_PE_=[x_ADP_ τ_ADP_ + (1-x_ADP_) τ_ADPꞏPi_]^−1^*, where *x_ADP_* is the average ADP-actin fraction of the filament, and *τ_ADPꞏPi_* (respectively, *τ_ADP_*) is the average time its takes to transform an ADPꞏP_i_-actin (respectively, ADP-actin) subunit at the cluster boundary into a cofilin-bound ADP-actin subunit. The fixed rate constants *k_on_* and *k_off_* for the growth of cofilin clusters on ADP-actin (Fig. S1a) determine *τ_ADP_*. Based on our results (Fig. 2g,h), we consider that cofilin binds with the same rate constants k_on_ and k_off_ to any ADP-actin subunit, regardless of the nucleotide state of the next subunit. The two following models differ in the way *τ_ADPꞏPi_* is computed.

Model 1 (Fig. 4c) considers that adding a cofilin when the next site is ADPꞏP_i_-actin requires the P_i_ to depart, at the enhanced rate *k ^model1^* , and to then add a cofilin on the newly ADP-actin site. This leads to *τ_ADPꞏPi_=1/ k ^model1^ + τ_ADP_*. The only unknown (free) parameter is *k ^model1^*.

Model 2 (Fig. 4d) considers that cofilin binds to the ADP*ꞏ*P_i_-actin subunit at the cluster boundary, with a dissociation constant *K_d_^model2^*, and that the P_i_ is released from the cofilin-bound ADP*ꞏ*P_i_-actin at rate *k ^model2^*. This takes place at a rate *τ ^−1^= k ^model2^ [cofilin]/([cofilin]+ K_d_ ^model2^).* There are two unknown (free) parameters, *K_d_^model2^* and *k ^model2^*

## SUPPLEMENTARY FIGURES

**Figure S1:**
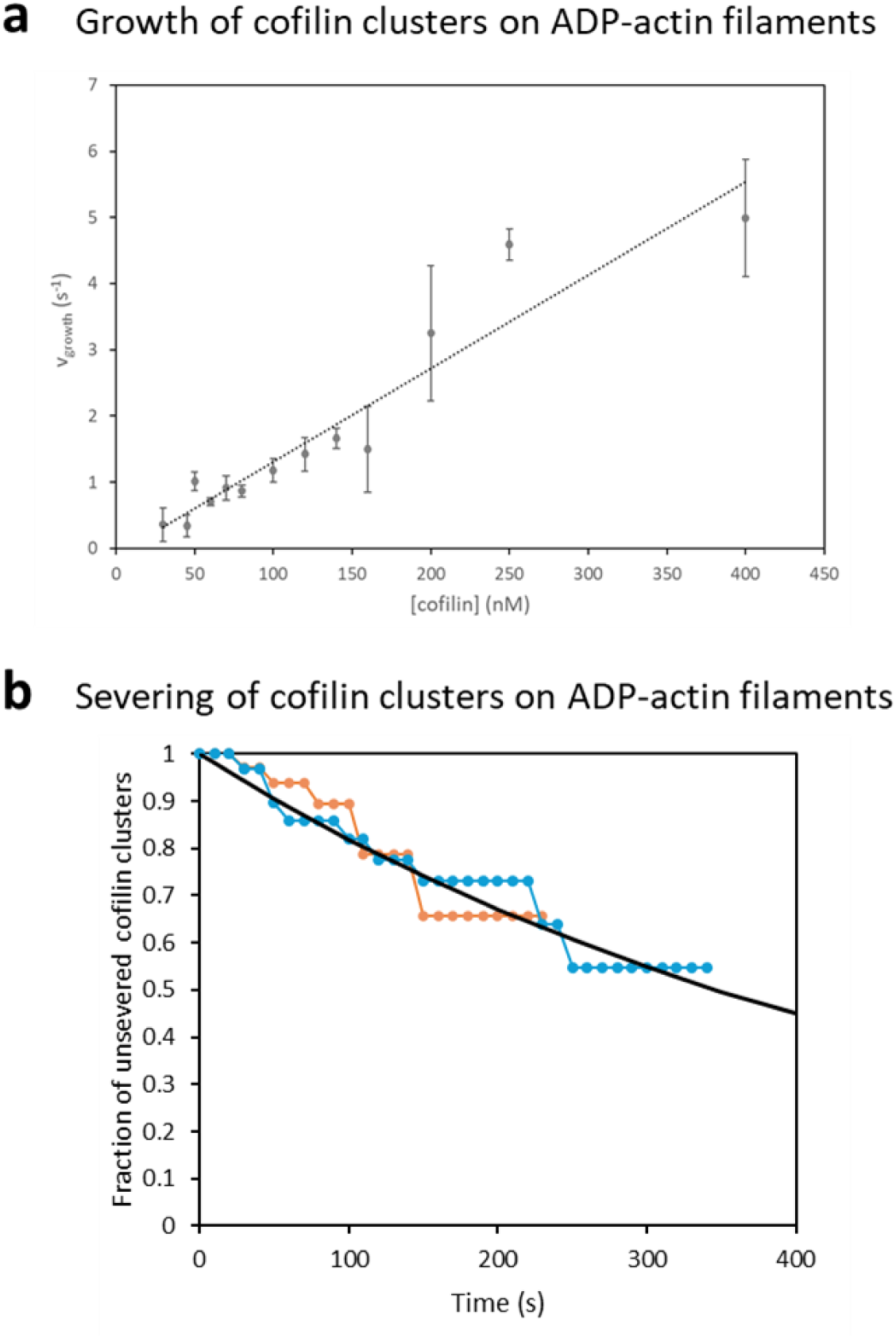
cluster growth and severing on ADP-actin filaments. **a.** Growth of cofilin clusters on ADP-actin filaments (both directions averaged together, as cluster growth is symmetrical, with v_BE_/v_PE_=1.08±0.11 averaged over 13 different cofilin concentrations), i.e. filaments that were aged for at least 15 minutes before being exposed to cofilin. Each point is the average over 5-188 different clusters. Bars indicate standard deviations. The dotted line is a linear fit, yielding k_on_=14 µM^−1^s^−1^ and k_off_ = 0.1 s^−1^. **b.** Severing at cofilin cluster boundaries on ADP-actin filaments. Data points are from two independent experiments, monitoring the severing of 36 (orange) and 37 clusters (blue) over time. The black line is an exponential fit, yielding a severing rate k_sev_=2ꞏ10^−3^ s^−1^ per cluster. We found that 84(±4)% (n=81) of severing events occurred at the cluster boundary toward the PE side of the filament.

**Figure S2:**
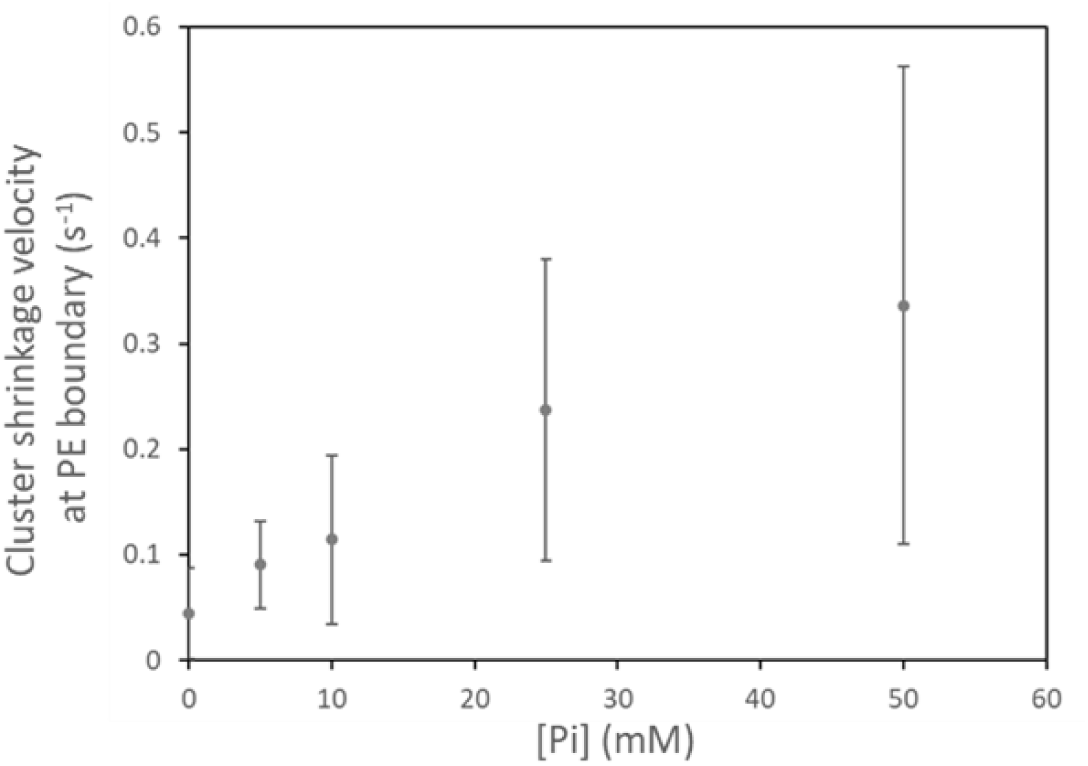
Shrinkage of cofilin clusters as a function of P_i_ concentration. Cofilin clusters formed on actin filaments were monitored while being exposed to buffer supplemented with P_i_, without cofilin (as in Fig. 2a-c). The velocity of the cluster boundary located toward the PE was measured. Each point is the average velocity for 15-34 clusters, from a series of experiments done on the same day, in the same microfluidics chamber. The clusters were shrinking, so the velocities can be viewed as negative velocities. Bars are standard deviations.

**Figure S3:**
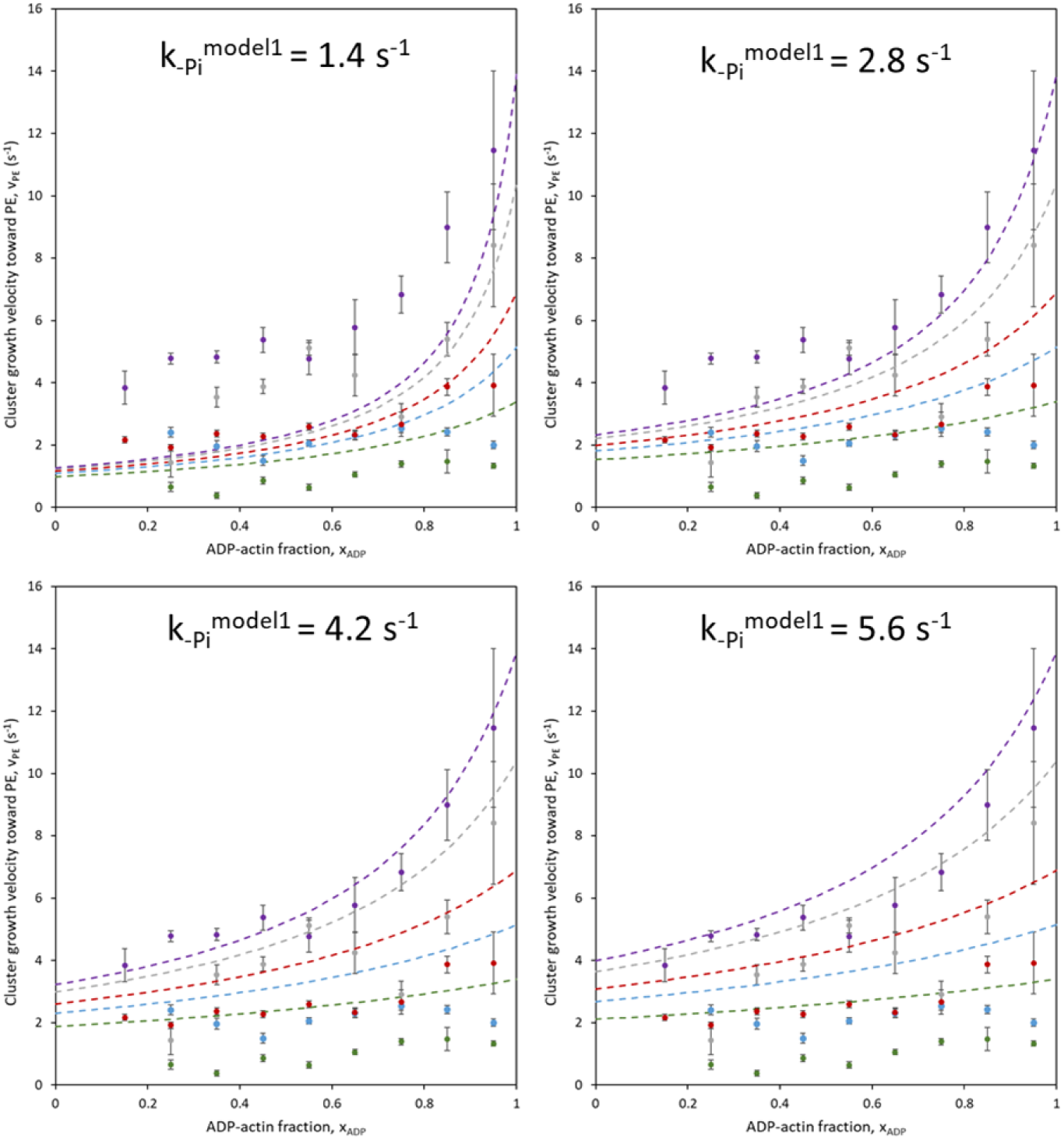
Examples of cluster growth velocities, computed using model1 with different values for the enhanced P_i_ release rate k_-Pi_^model1^.

**Figure S4:**
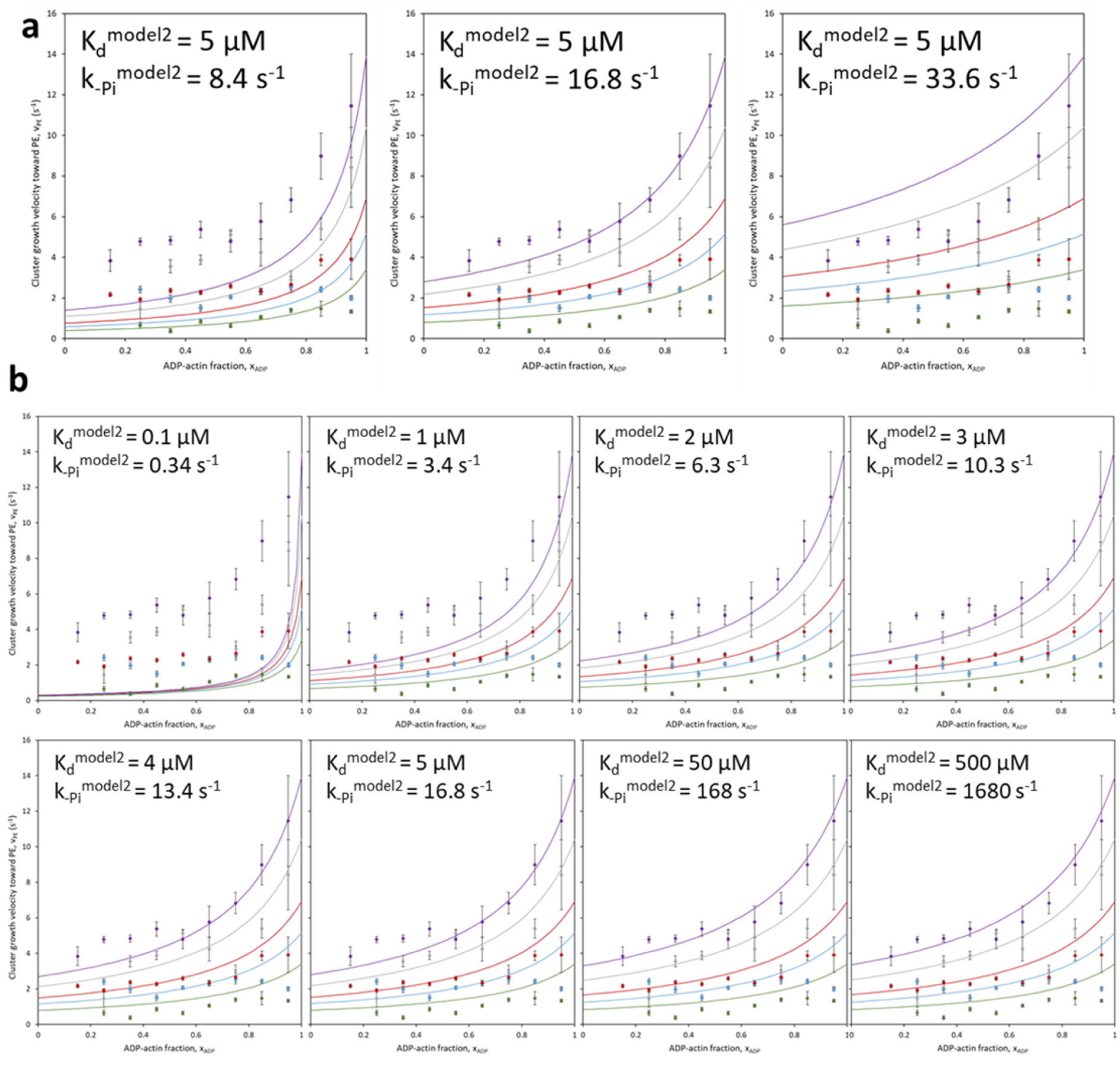
Examples of cluster growth velocities, computed using model2 with different parameters. **a.** Sets of curves computed with the same K_D_^model2^= 5 µM and different values for k_-Pi_^model2^. **b.** Sets of curves all computed with the same ratio K_D_^model2^ / k_-Pi_^model2^ ∼0.3 µMꞏs^−1^, and different values for the two parameters. The experimental data (same as in Fig. 4c,d) is shown as a reference.

## Notes

### Competing Interest Statement

The authors have declared no competing interest.

